# Diversity and metagenome analysis of a hydrocarbon-degrading bacterial consortium from asphalt lakes located in Wietze, Germany

**DOI:** 10.1101/2021.03.25.436929

**Authors:** Michael O. Eze, Grant C. Hose, Simon C. George, Rolf Daniel

## Abstract

The pollution of terrestrial and aquatic environments by petroleum contaminants, especially diesel fuel, is a persistent environmental threat requiring cost-effective and environmentally sensitive remediation approaches. Bioremediation is one such approach, but is dependent on the availability of microorganisms with the necessary metabolic abilities and environmental adaptability. The aim of this study was to examine the microbial community in a petroleum contaminated site, and isolate organisms potentially able to degrade hydrocarbons. Through successive enrichment of soil microorganisms from samples of an historic petroleum contaminated site in Wietze, Germany, we isolated a bacterial consortium using diesel fuel hydrocarbons as sole carbon and energy source. The 16S rRNA gene analysis revealed the dominance of *Alphaproteobacteria*. We further reconstructed a total of 18 genomes from both the original soil sample and the isolated consortium. The analysis of both the metagenome of the consortium and the reconstructed metagenome-assembled genomes show that the most abundant bacterial genus in the consortium, *Acidocella*, possess many of the genes required for the degradation of diesel fuel aromatic hydrocarbons, which are often the most toxic component. This can explain why this genus proliferated in all the enrichment cultures. Therefore, this study reveals that the microbial consortium isolated in this study and its dominant genus, *Acidocella*, could potentially serve as an effective inoculum for the bioremediation of sites polluted with diesel fuel or other organic contaminants.

## INTRODUCTION

Petroleum pollution is a recurring environmental threat resulting from oil and gas exploration, production, transport and storage (Eze and George 2020). Spills have occurred in terrestrial as well as aquatic environments, and they are often caused by human error, corrosion and equipment failure (Dalton and Jin 2010; Errington et al. 2018; Hassler 2016; Hong et al. 2010). This is a major threat to both the environment and human health, due to the phytotoxicity and carcinogenicity of petroleum hydrocarbons.

In view of the diversity of pollutants, a range of *ex situ* and *in situ* bioremediation techniques have been developed (Azubuike et al. 2016). *Ex situ* techniques involve the excavation and off-site treatment of contaminated soils or water, while *in situ* strategies involve on-site treatment of contaminants. As a result, *ex situ* techniques are often more expensive than *in situ* techniques owing to the additional costs associated with contaminant excavation and relocation (USEPA 2000). The United States Environmental Protection Agency indicated that implementing *in situ* degradation will result in cost savings of 50 to 80% over traditional methods such as excavation and landfill incineration (USEPA 2001). Moreover, *ex situ* methods are environmentally problematic as they alter the soil matrix and associated microbiomes.

The success of any oil spill remediation approach depends on environmental conditions such as temperature, pH and nutritional constraints in contaminated sites (Joner et al. 2002; Kleinsteuber et al. 2006; Leahy and Colwell 1990; Rohrbacher and St-Arnaud 2016). The hydrophobic nature of petroleum hydrocarbons limits their availability to biodegradation. Hence, the presence of microorganisms with the metabolic capability to degrade petroleum and the ability to adapt to a range of environmental conditions is a crucial factor (Das and Chandran 2011). Organisms capable of degrading diesel fuel and other organic contaminants are diverse and present in many natural habitats, including extreme ones (Gemmell and Knowles 2000; Hara and Uchiyama 2013; Lohi et al. 2008; Nie et al. 2014; Stapleton et al. 1998). Microorganisms from polluted environments hold the key to unlocking most of the challenges associated with bioremediation (Eze et al. 2020; Liang et al. 2019; Liang et al. 2016). One such environment is the heavily polluted oil field in Wietze, Germany.

Wietze is an important historical site of crude-oil production. In Germany, pre-industrial oil production started in the 17^th^ century, followed by industrial oil extraction beginning in 1859 (Craig et al. 2018). Between 1900 and 1920, Wietze was the most productive oil field in Germany, with almost 80% of German oil produced there. Oil production in Wietze was discontinued in 1963, but the former oil field continues to witness considerable amounts of oil seepage, with several heavily polluted sites, contaminated ponds, and organic debris from surrounding plants (Figure 1). Therefore, it is an ideal site for obtaining microorganisms with the potential for bioremediation of petroleum hydrocarbons. Samples investigated in this study were taken from three sites around a small asphalt pond (Figure 1).

**Figure 1.**
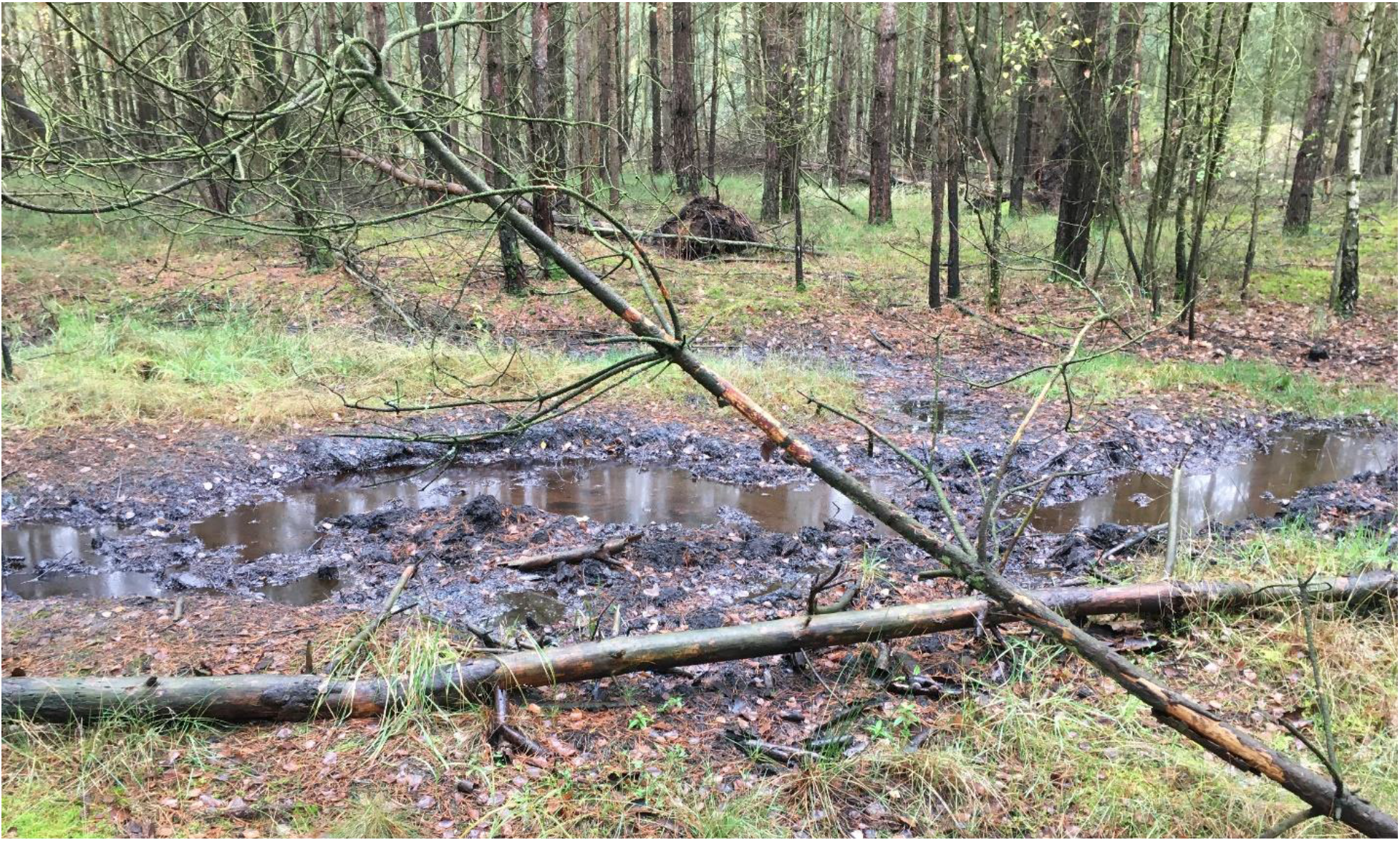
Sampling site in Wietze, Germany (52°39’0’’N, 09°50’0’’E).

Due to the so-called uncultivability of many environmental microorganisms (Steen et al. 2019), several studies have concentrated on remediation by indigenous microorganisms (Kumar and Gopal 2015; Sarkar et al. 2016). More recent studies have shown that the inoculation of carefully cultivated hydrocarbon-degrading bacterial consortia or isolates enhances the effectiveness of various remediation techniques (Atashgahi et al. 2018; Garrido-Sanz et al. 2019). Therefore, it is important to discover novel microbes capable of degrading petroleum hydrocarbons either as single isolates, consortia, or synergistically with plants. The aim of this study was to investigate the diversity and genomic potential of bacterial consortia derived from a hydrocarbon contaminated asphalt lake in Wietze, Germany. We also aimed to reconstruct metagenome-assembled genomes, and to examine the potential of the reconstructed genomes for bioremediation of diesel fuel contaminated sites.

## MATERIALS AND METHODS

### Soil sampling

Topsoil samples (10 g each) and water samples (approximately 50 mL each) were taken in November 2019 from three heavily polluted sites located at the historical oil field in Wietze (52°39’0’’N, 09°50’0’’E), Germany. In addition, two reference samples were taken from nearby unpolluted soils. Samples were placed into 50 mL Eppendorf conical tubes. The samples were transported to the laboratory on ice.

### Enrichment cultures and growth conditions

Approximately 1 g of each of the crude oil-polluted soil samples was added to Erlenmeyer flasks (300 mL) containing 100 mL of a liquid mineral medium (MM) composed of KH_2_PO_4_ (0.5 g/L), NaCl (0.5 g/L), and NH_4_Cl (0.5 g/L). Sterile-filtered trace elements (1 mL/L) (Atlas 2010), vitamin solution (1 mL/L) (Atlas 2010) and MgSO_4_.7H_2_O (5 mL/L of a 100 mg/mL solution) were added to the MM, post MM-autoclaving. One mL of sterile-filtered diesel fuel (C_10_-C_25_) was added to each flask as the sole carbon and energy source. The cultures were grown at 30°C with shaking at 110 rpm (INFORS HT shaker, model CH-4103, Infors AG, Bottmingen, Switzerland) and subcultured every five days. After three successive subculturing steps, 30 mL aliquots (OD_600_, 0.635) were centrifuged for 10 min at 4,000 × *g*.

### DNA Extraction

Microbial cells from approximately 30 mL of the enrichment cultures and water samples were harvested by centrifugation at 4,000 x *g* for 10 min. The supernatant was subsequently discarded. DNA from the cell pellets and 100 mg of each of the original samples were extracted using the PowerSoil^®^ DNA Extraction kit as recommended by the manufacturer (Qiagen, Hilden, Germany). DNA from one of the original soil samples and one of the three final enrichments (S3S and S3E3 respectively, Supplementary Figure S1) were used for metagenome studies.

### Sequencing of bacterial 16S rRNA genes

Bacterial 16S rRNA genes were amplified using the forward primer S-D-Bact-0341-b-S-17 (5′-CCT ACG GGN GGC WGC AG-3′) and the reverse primer S-D-Bact-0785-a-A-21 (5′-GAC TAC HVG GGT ATC TAA TCC-3′) (Klindworth et al. 2013) containing adapters for Illumina MiSeq sequencing. The PCR reaction (25 μL final volume) contained 5 μL of five-fold Phusion HF buffer, 200 μM of each of the four deoxynucleoside triphosphates, 4 μM of each primer, 1 U of Phusion high fidelity DNA polymerase (Thermo Scientific, Waltham, MA, USA), and approximately 50 ng of the extracted DNA as the template. Negative controls were performed using the reaction mixture without a template. The following thermal cycling scheme was used: initial denaturation at 98 °C for 30 s, 30 cycles of denaturation at 98 °C for 15 s, annealing at 53 °C for 30 s, followed by extension at 72 °C for 30 s. The final extension was carried out at 72 °C for 2 min. The PCR products that were obtained were controlled for appropriate size, and then purified using the MagSi-NGS Plus kit according to the manufacturer’s protocol (Steinbrenner Laborsysteme GmbH, Germany). Quantification of the PCR products was performed using the Quant-iT dsDNA HS assay kit and a Qubit fluorometer, as recommended by the manufacturer (Thermo Scientific). The DNA samples were barcoded using the Nextera XT-Index kit (Illumina, San Diego, USA) and the Kapa HIFI Hot Start polymerase (Kapa Biosystems, USA). Sequencing was performed at the Göttingen Genomics Laboratory using an Illumina MiSeq Sequencing platform (paired-end 2 × 300 bp) and the MiSeq reagent kit v3, as recommended by the manufacturer (Illumina).

### Processing of the 16S rRNA gene data

Trimmomatic version 0.39 (Bolger et al. 2014) was initially used to truncate low-quality reads if quality dropped below 12 in a sliding window of 4 bp. Datasets were subsequently processed with Usearch version 11.0.667 (Edgar 2010) as described in Wemheuer et al. (2020). In brief, paired-end reads were merged and quality-filtered. Filtering included the removal of low-quality reads and reads shorter than 200 bp. Processed sequences of all samples were joined, dereplicated and clustered in zero-radius operational taxonomic units (zOTUs) using the UNOISE algorithm implemented in Usearch. A *de novo* chimera removal was included in the clustering step. Afterwards, zOTU sequences were taxonomically classified using the SINTAX algorithm against the SILVA database (SILVA SSURef 138 NR99). All non-bacterial zOTUs were removed based on taxonomic classification. Subsequently, processed sequences were mapped on final zOTU sequences to calculate the distribution and abundance of each OTU in every sample.

### Metagenome sequencing, assembly and analysis

Sequencing libraries were generated from environmental DNA. These were barcoded using the Nextera XT-Index kit (Illumina, San Diego, USA) and the Kapa HIFI Hot Start polymerase (Kapa Biosystems, Wilmington, USA). Sequencing was performed by employing an Illumina HiSeq 2500 system and the HiSeq Rapid SBS kit V2 (2×250 bp) as recommended by the manufacturer (Illumina). Metagenomic reads were further processed as described previously (Eze et al. 2020). In brief, reads were processed with the Trimmomatic tool version 0.39 (Bolger et al. 2014) and assembled using metaSPAdes version 3.13.2 (Bankevich et al. 2012). Coverage information for each scaffold was determined using Bowtie2 version 2.3.2 (Langmead and Salzberg 2012) and SAMtools version 1.7 (Li et al. 2009). Metagenome-assembled genomes (MAGs) were reconstructed with MetaBAT version 2.12.1 (Kang et al. 2015). MAG quality was determined using CheckM version 1.0.13 (Parks et al. 2015). Only MAGs with a completeness minus contamination of more than 50% and a contamination rate of less than 7% were considered for further analysis. MAGs were classified taxonomically using GTDB-Tk version 1.0.2 and the Genome Taxonomy Database (release 86) (Chaumeil et al. 2019; Parks et al. 2019). Coding DNA sequences (CDSs) were identified with prodigal version 2.6.3 (Hyatt et al. 2010). Functional annotation was performed with diamond version v0.9.29 (Buchfink et al. 2015) and the KEGG database (October release 2018) (Kanehisa and Goto 2000).

### Data analysis

Data analysis was performed in R (RCoreTeam 2018). Richness, diversity, evenness, and coverage based on the Chao1 richness estimator were estimated in R using the vegan package (RCoreTeam 2018). In addition, richness was estimated using the Michaelis-Menten equation in R with the drc package (RCoreTeam 2018). Prior to alpha diversity analysis, the zOTU table was rarefied to 12,924 per sample. Beta-diversity was calculated in R using the vegan package. Non-metric multidimensional scaling plots were generated based on Bray-Curtis dissimilarities. Dissimilarities were calculated based on the raw zOTU table.

## RESULTS

### Bacterial diversity of the sampling sites and the diesel-degrading cultures

The 16S rRNA gene amplicon sequencing resulted in 242,025 16S rRNA gene sequences across all samples (36,441–10,309 reads per sample, average 22,002 per sample). Clustering resulted in a total of 6,453 zOTUs (average: 587) ranging from 225 to 813 zOTUs per sample. The highest bacterial richness and diversity were observed in the reference samples, the lowest in the enrichment samples. Calculated coverage values indicate that the majority of the bacterial diversity (>80.9%, see Supplementary Table S1) was recovered by the surveying effort.

Non-metric multidimensional scaling revealed clear differences between the microbial community composition of the polluted soil and water samples, enrichment cultures, and reference unpolluted soil samples (Figure 2).

**Figure 2.**
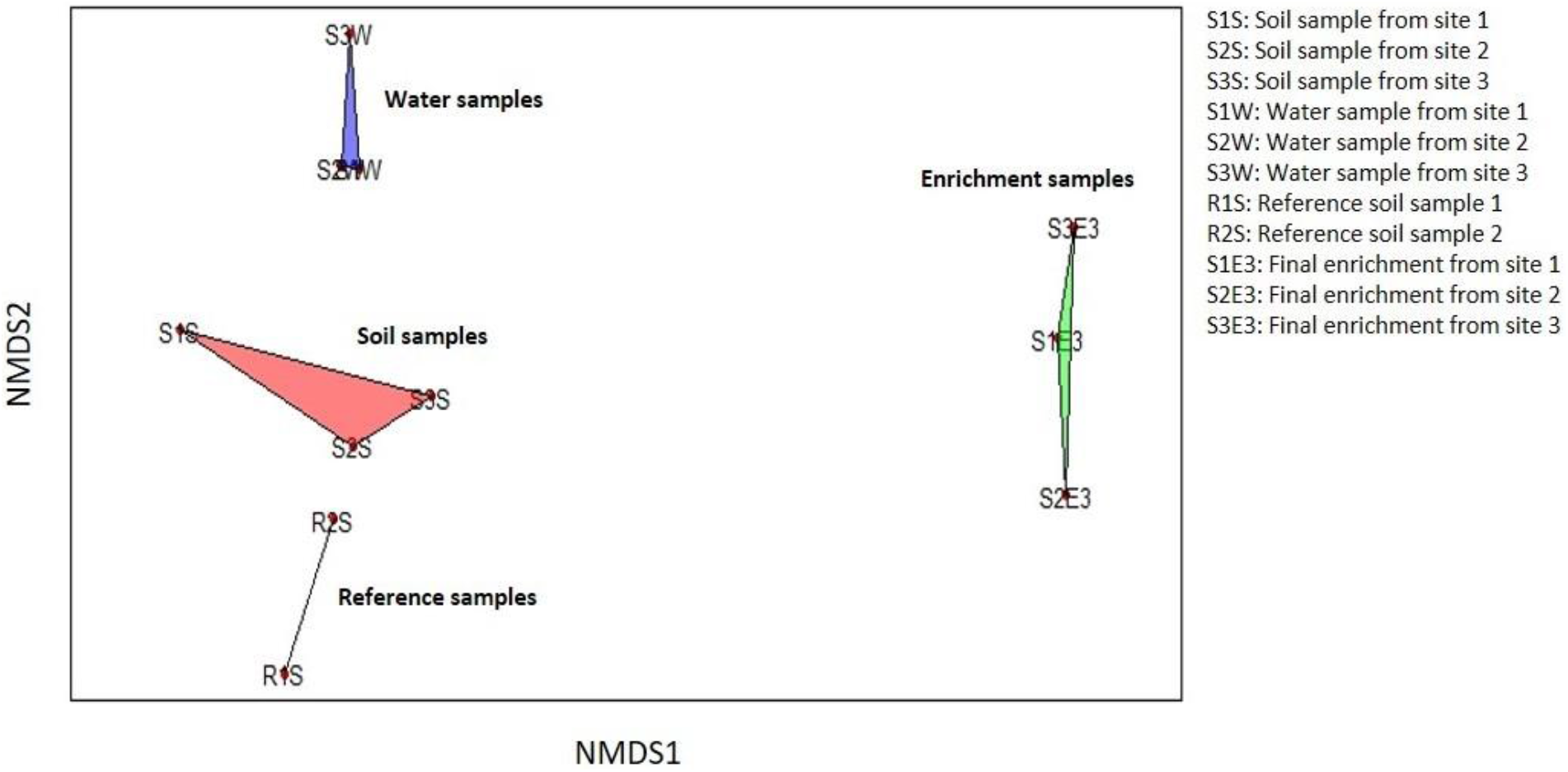
Non-metric multidimensional scaling (NMDS) ordination plot showing differences in microbial community compositions of the water, soil, enrichment, and reference unpolluted soil samples based on community composition at the genus level.

The relative abundances at the bacterial class level (Figure 3a) showed the dominance of *Gammaproteobacteria* in the polluted water sample (90.6%), followed by *Alphaproteobacteria* (3.2%). The polluted soil samples contain similar relative abundances for *Gammaproteobacteria*, *Alphaproteobacteria* and *Acidobacteriae* (26.4%, 21.4% and 19.1%, respectively). The enrichment cultures are dominated by members of the *Alphaproteobacteria*, with a relative abundance of 75.8%. Other bacterial classes present in the enrichment culture include *Gammaproteobacteria* and *Acidobacteriae* (15.4% and 8.6%, respectively). A higher diversity and richness (Supplementary Table S1) was recorded in the unpolluted reference sample in which *Actinobacteria* (17.0%), *Alphaproteobacteria* (14.6%), *Acidobacteriae* (13.5%), and *Bacteroidia* (10.1%) are dominant. Other less abundant classes include *Phycisphaerae* and *Verrucomicrobiae*. At genus level, *Acidocella* are dominant in all the enrichment cultures from the three sites (87.4% to 75.4%). Other genera present in the enrichment cultures include *Acidobacterium* and *Paraburkholderia* (Figure 3b).

**Figure 3.**
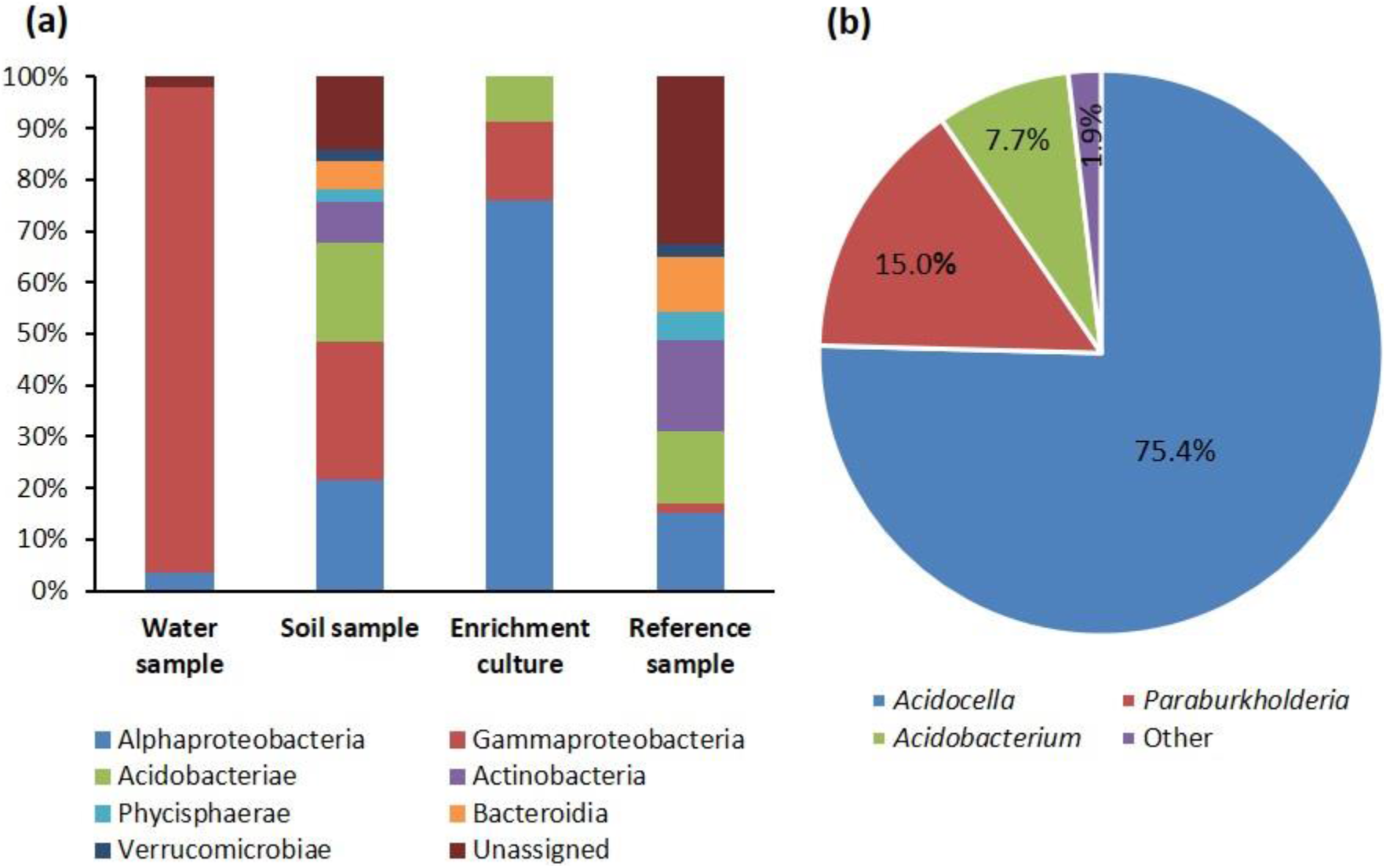
(a) Bacterial community composition in selected water, soil, enrichment, and unpolluted reference samples. (b) relative abundance of the enrichment culture at the genus level. Only taxa with a relative abundance of >1% across all samples are presented. For details on relative abundances and 16S rRNA gene amplicon data, see Supplementary Figure S1.

### Identification of aliphatic and aromatic hydrocarbon-degrading coding DNA sequences

Functional analysis of the metagenome derived from the microbial diesel enrichment revealed the presence of 42 potential enzymatic classes represented by 186 coding DNA sequences (CDSs) involved in the degradation of aliphatic and aromatic hydrocarbons (Figure 4).

**Figure 4.**
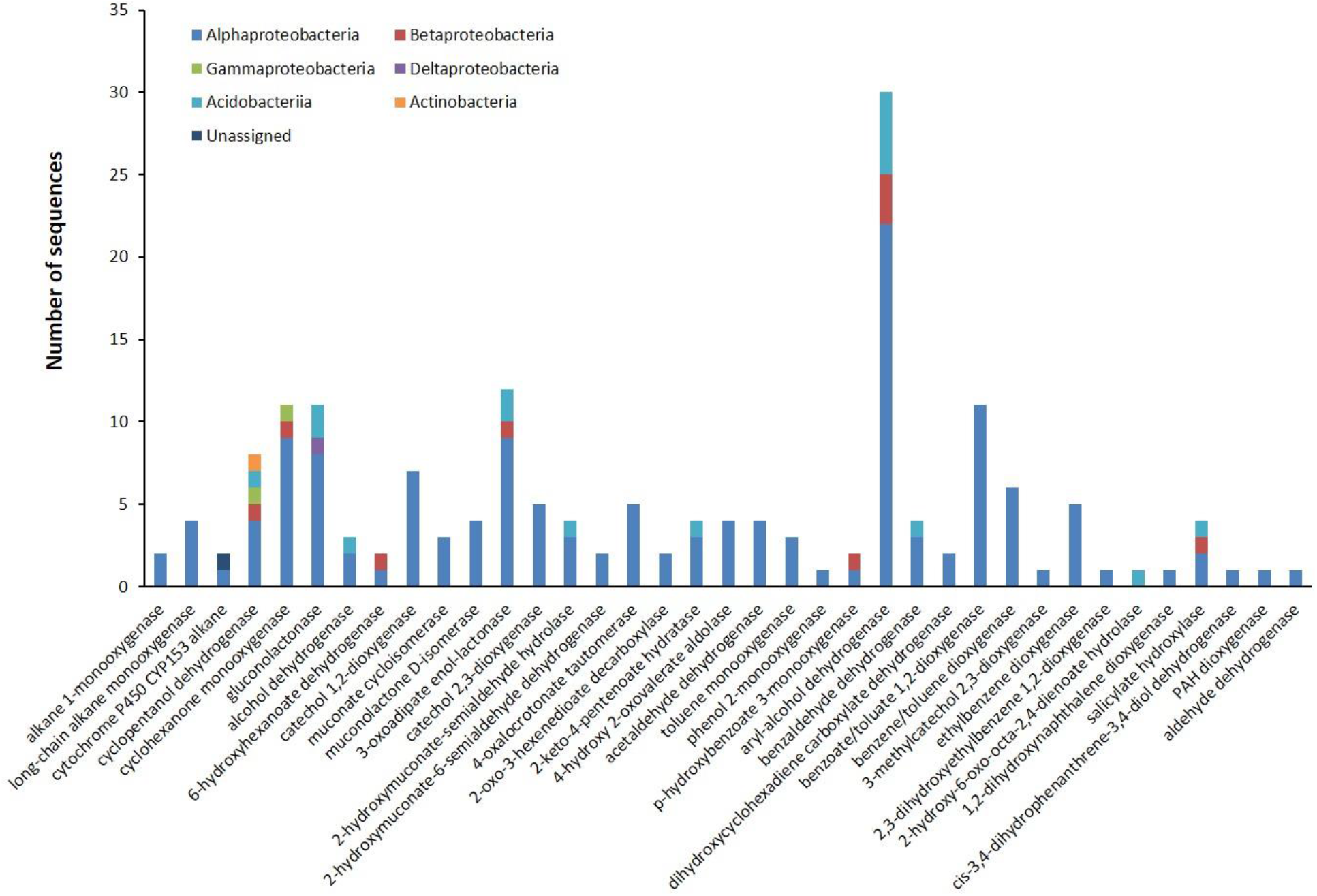
The number of sequences associated with specific hydrocarbon-degrading enzymes in each taxonomic group. The analysis was based on the metagenome of the S3E3 enrichment culture.

The enzymes considered as responsible for the degradation of aliphatic hydrocarbons included alkane 1-monooxygenase, long-chain alkane monooxygenase, cytochrome P450 CYP153 alkane hydroxylase, cyclopentanol dehydrogenase, cyclohexanone monooxygenase, gluconolactonase, alcohol dehydrogenase, and 6-hydroxyhexanoate dehydrogenase. Forty-three CDSs were detected that are considered to play a role in aliphatic hydrocarbon degradation. The majority of the genes that putatively code for aliphatic hydrocarbon degradation are involved in cycloalkane degradation. These include the *cpnA*, *chnB, gnl*, *adh* and *chnD* genes, which are involved in the Baeyer-Villiger oxidation reactions (Figure 5).

**Figure 5.**
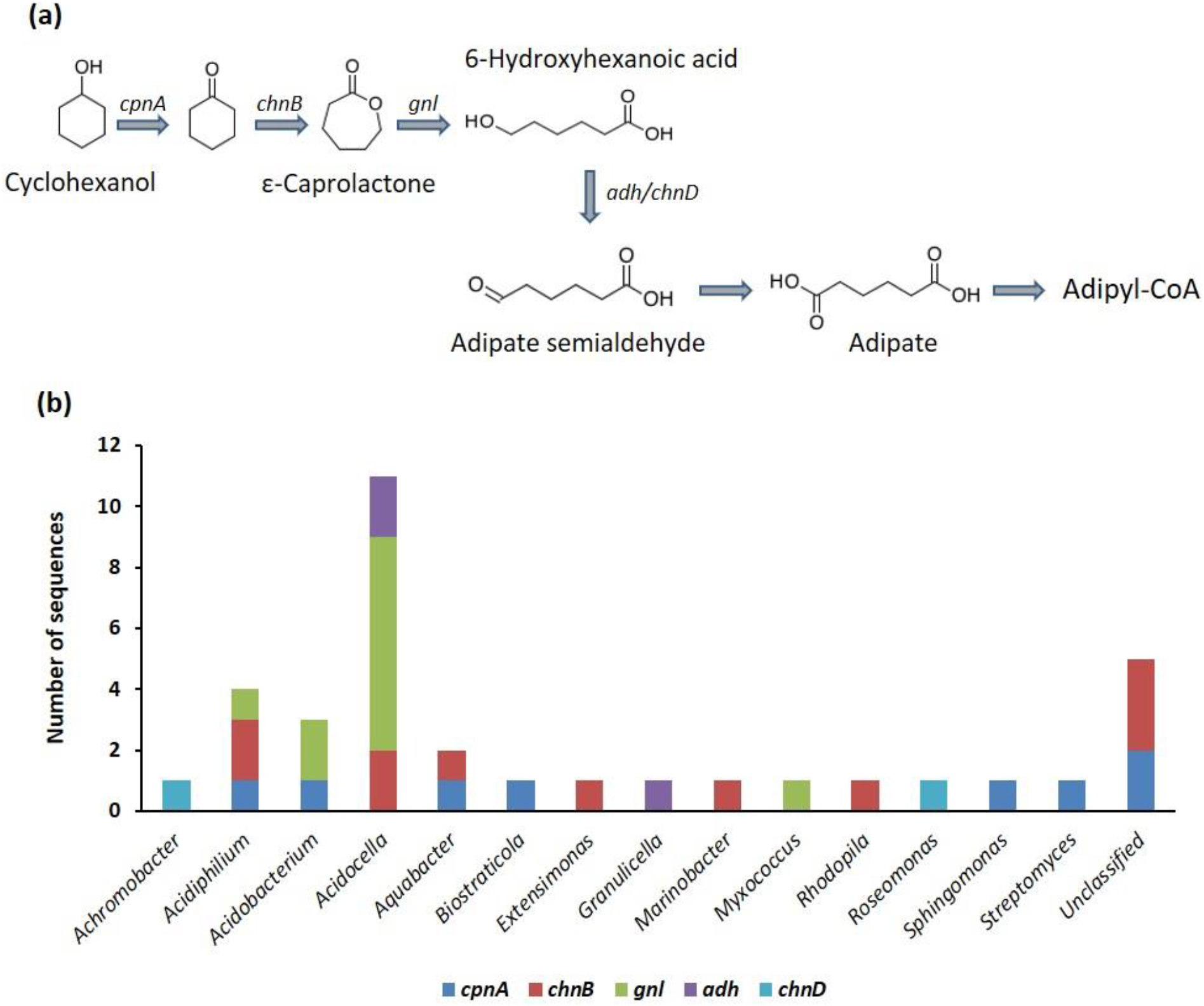
(a) Ring cleavage via the Baeyer-Villiger oxidation pathway for the degradation of cycloalkanes, and (b) genus assignment of the putative genes involved in the Baeyer-Villiger oxidation pathway identified in the diesel-degrading consortium.

The degradation of aromatic hydrocarbons occurs through a series of reactions involving oxidation, hydroxylation, dehydrogenation and ring cleavage. Out of the 186 CDSs putatively linked to diesel degradation, 143 CDSs are potentially involved in aromatic hydrocarbon degradation. Among the 48 CDSs belonging to the aromatic ring dioxygenases, eleven were annotated as benzoate/toluate 1,2-dioxygenase, six as biphenyl 2,3-dioxygenase, six as benzene/toluene/chlorobenzene dioxygenase, five as ethylbenzene dioxygenase, and three as naphthalene 1,2-dioxygenase (Table 1).

**Table 1.** Key monooxygenases and dioxygenases involved in the activation and ring cleavage of aromatic hydrocarbons in the diesel-degrading consortium.

| Genes | Enzyme | Function | No. of CDSs |
| --- | --- | --- | --- |
| *tmoCF* | Toluene monooxygenase | Activation | 3 |
| *pobA* | p-Hydroxybenzoate 3-monooxygenase | Activation | 2 |
| *todABC1C2* | Benzene/toluene/chlorobenzene dioxygenase | Activation | 6 |
| *etbAaAbAc* | Ethylbenzene dioxygenase | Activation | 5 |
| *benABC* | Benzoate/toluate 1,2-dioxygenase | Activation | 11 |
| *bphA* | Biphenyl 2,3-dioxygenase | Activation | 6 |
| *nahAb* | Naphthalene 1,2-dioxygenase | Activation (PAHs) | 3 |
| *nahC* | 1,2-Dihydroxynaphthalene dioxygenase | Activation (PAHs) | 1 |
| *nidA* | PAH dioxygenase | Activation (PAHs) | 1 |
| *catA* | Catechol 1,2-dioxygenase | Ortho-cleavage | 7 |
| *dmpB* | Catechol 2,3-dioxygenase | Meta-cleavage | 5 |
| *todE* | 3-Methylcatechol 2,3-dioxygenase | Meta-cleavage | 1 |
| *etbC* | 2,3-Dihydroxyethylbenzene 1,2-dioxygenase | Meta-cleavage | 1 |
| *bphC* | Biphenyl-2,3-diol 1,2-dioxygenase | Meta-cleavage | 1 |

### Reconstruction of metagenome-assembled genomes

We were able to reconstruct fifteen nearly complete genomes from the whole-metagenome sequence of the original soil samples, and three nearly complete genomes from the enrichment culture (Supplementary Table S2). Quality analysis of the MAGs showed that the average completeness and contamination level for the MAGs were 85% and 2% respectively (Supplementary Table S3). The majority of the metagenome-assembled genomes (MAGs) were classified as belonging to the *Gammaproteobacteria* (8 MAGs), followed by *Alphaproteobacteria* (4 MAGs), *Acidobacteriae* (3 MAGs), *Actinobacteria* (2 MAGs) and *Caldisericia* (1 MAG). The three metagenome-assembled genomes from the enrichment culture were classified as *Acidocella aminolytica*, *Acidobacterium capsulatum*, and *Acidocella* sp., with a completeness of 72.4%, 99.8% and 100%, respectively.

A comparison of the three nearly complete genomes reconstructed from the metagenome of the enrichment culture shows that the genes encoding enzymes involved in the activation and degradation of petroleum hydrocarbons are more abundant in *Acidocella* than in *Acidobacterium* (Supplementary Table S2). For example, while the two MAGs classified as *Acidocella* contain an average of 18 CDSs involved in aromatic ring activation, *Acidobacterium* had only 7 CDSs encoding for the activation of aromatic hydrocarbons. Key enzymes that are encoded by the reconstructed MAGs belonging to *Acidocella* but are missing in those belonging to *Acidobacterium* include long-chain alkane monooxygenase, cyclohexanone monooxygenase, ethylbenzene dioxygenase, and benzoate/toluate 1,2-dioxygenase.

Further comparisons performed between the MAGs assembled from the metagenome data of the enrichment culture and those obtained from a previous study of a crude oil bore hole (Eze et al. 2020) revealed that the *Acidocella* MAGs obtained from this study exhibit a higher abundance of genes that putatively encode the degradation of cycloalkanes. For example, in the 36 MAGs from Eze et al. (2020), genes that encode for cyclopentanol dehydrogenase (*cpnA*) and cyclohexanone monooxygenase (*chnB*) were present in 16 and 11 MAGs, respectively. In this study, MAGs reconstructed from both the enrichment culture and the original soil samples were rich in genes that encode these enzymes with more than 6 CDSs per gene in some MAGs. The reconstructed MAGs were also found to be richer in CDSs that encode for aromatic degradation that those in the previous study. For example, aryl alcohol dehydrogenase, an enzyme vital for the degradation of aromatic hydrocarbons was missing in all of the 36 assembled MAGs from the crude oil bore hole study (Eze et al. 2020). Potential genes encoding the enzyme were present in two of the three MAGs from the enrichment culture of this study.

## DISCUSSION

The successive enrichment of the different experimental samples using diesel fuel resulted in the dominance of *Alphaproteobacteria*. The dominance of *Alphaproteobacteria* in the bacterial communities, especially *Acidocella* and *Paraburkholderia*, indicates the tolerance of these genera to high concentrations of petroleum hydrocarbons and their potential degradative capacity for organic contaminants. The taxa that are abundant in the polluted water and soil, and in the enrichment cultures, were also associated with hydrocarbon pollution in other locations (Lee et al. 2019; Röling et al. 2006; Stapleton et al. 1998). The biodegradative ability of these taxa and their tolerance to heavy metals (Giovanella et al. 2020) indicate that they are potentially suitable for the remediation of multiple contaminants such as hydrocarbon-polluted acidic mine sites.

Diesel fuel contains aliphatic and aromatic hydrocarbons. The aliphatic hydrocarbon fraction is predominantly composed of normal-, iso- and cyclo-alkanes, while the aromatic hydrocarbon fraction is composed primarily of alkylbenzenes, naphthalene, alkylnaphthalenes, biphenyl and alkylbiphenyls (Woolfenden et al. 2011). The degradation of n-alkanes is primarily carried out by alkane 1-monooxygenase (*alkB*), cytochrome P450 CYP153 alkane hydroxylase (*CYP153*) and long-chain alkane monooxygenases (*ladA*) genes, and their roles in the degradation of *n-*alkanes and iso-alkanes have been extensively studied (Ji et al. 2013; Li et al. 2008; van Beilen et al. 2006). The degradation of *n*-alkanes and iso-alkanes by the consortium is indicated by the presence of potential *alkB*, *CYP153* and *ladA* genes. The low number of the corresponding gene sequences (eight) can be explained by the taxonomic composition of the consortium. Previous studies have shown that n-alkane degrading genes are often associated with *Betaproteobacteria* and *Gammaproteobacteria* especially the *Pseudomonas* genus (Garrido-Sanz et al. 2019; Liu et al. 2014; Shao and Wang 2013; van Beilen et al. 2001; van Beilen et al. 1994). In our study, the diesel-degrading consortium in the enrichment cultures was dominated by *Alphaproteobacteria* (Figures 2 and 3). Thus, the majority of CDSs in our metagenome consortium belong to the *Alphaproteobacteria*, especially the *Acidocella* genus and not to *Pseudomonas*.

Of the genes that putatively code for aliphatic hydrocarbon degradation, the majority are involved in cycloalkane degradation. These enzymes include cyclopentanol dehydrogenase (*cpnA*), cyclohexanone monooxygenase (*chnB*), gluconolactonase (*gnl*), alcohol dehydrogenase (*adh*), and 6-hydroxyhexanoate dehydrogenase (*chnD*) (Bohren et al. 1989; Iwaki et al. 1999; Iwaki et al. 2002; Kanagasundaram and Scopes 1992). This is interesting since cycloalkanes are moderately resistant to biodegradation (Connan 1984). The degradation of cycloalkanes involves ring cleavage via Baeyer-Villiger oxidation (Perkel et al. 2018; Sheng et al. 2001), which requires an initial oxidation of cyclohexane to cyclohexanol by cyclohexane monooxygenase, and then a dehydrogenation reaction to cyclohexanone. This step is followed by another monooxygenase attack to form epsilon-caprolactone, followed by ring cleavage that is carried out by gluconolactonase (Figure 5a). All the genes involved in this degradation pathway are present in the metagenome of the enrichment culture, but a single taxon in the bacterial community that possess all the genes involved in this pathway was not detected (Figure 5b). This indicates a synergistic interaction of different bacterial genera in the degradation of recalcitrant hydrocarbons. The high number of *cpnA, chnB, gnl, adh* and *chnD* genes (35 CDSs) in the metagenome of the enrichment culture indicates the significant potential of the microbial community for the degradation of cycloalkanes present in diesel fuel.

The degradation of aromatic hydrocarbons requires initial activation by oxygenases resulting in the formation of oxygenated intermediates such as catechol (Atashgahi et al. 2018; Das and Chandran 2011; Peters et al. 2004). The bacterial consortium contains more genes that putatively encode dioxygenases than those that encode monooxygenases (Table 1). The genes that encode dioxygenases include the *todABC1C2*, *etbAaAbAc* and *benABCD* genes (Fong et al. 1996; Werlen et al. 1996; Zylstra and Gibson 1989). The higher abundance of genes encoding dioxygenases indicates that the activation of alkylbenzenes and phenolic compounds by the microbial consortium predominantly follows the dioxygenase pathway rather than the monooxygenase pathway.

The central metabolism of aromatic hydrocarbons that follows initial activation involves ortho- and meta-cleavage of catechol or methylcatechol (Benjamin et al. 1991; Ehrt et al. 1995; Hidalgo et al. 2020; Liang et al. 2019; Neidle et al. 1988; Peters et al. 2004; Rohrbacher and St-Arnaud 2016). Functional analysis reveals that genes encoding enzymes putatively involved in the central metabolism of aromatic hydrocarbons are present in the microbial community. The most abundant CDSs in our diesel-degrading community that are responsible for this reaction are catechol 1,2-dioxygenase and catechol 2,3-dioxygenase (7 and 5 CDSs, respectively) (Table 1). Other enzymes that are present include 3-oxoadipate enol-lactonase, muconolactone D-isomerase (a decarboxylating dehydrogenase), 4-oxalocrotonate tautomerase, and acetaldehyde dehydrogenase. Most of the corresponding genes are affiliated to *Alphaproteobacteria*.

Polycyclic aromatic hydrocarbons (PAHs) are more resistant to microbial attack than smaller aromatic hydrocarbons, and when biodegradation is possible, this often proceeds through oxidation and ring cleavage by dioxygenases (Sipilä et al. 2008). The metagenome contains genes that encode enzymes putatively involved in the degradation of PAHs and other recalcitrant hydrocarbons, such as biphenyl and alkylbiphenyls. These enzymes include naphthalene 1,2-dioxygenase (*nahAb*) and 1,2-dihydroxynaphthalene dioxygenase (*nahC*) for naphthalene and alkylnaphthalenes (Peng et al. 2008), biphenyl 2,3-dioxygenase (*bphA*) for biphenyl and alkylbiphenyls, and PAH dioxygenase (*nidA*) for phenanthrene, alkylphenanthrenes, and other high molecular weight PAHs (Iwasaki et al. 2006; Robrock et al. 2011) (Table 1). Since crude oil and oil spills often contain significant amount of polycyclic aromatic hydrocarbons such as naphthalene, alkylnaphthalenes, phenanthrene and alkylphenanthrenes (Ahmed and George 2004; Eze and George 2020), the presence of putative genes encoding PAH dioxygenases in the metagenome of the consortium indicates the potential of the consortium for the remediation and reclamation of petroleum-contaminated soils.

Interestingly, the majority of previous studies on microbially-enhanced rhizoremediation of petroleum hydrocarbons have focused on *Pseudomonas* (de Lima-Morales et al. 2015; Di Martino et al. 2012), *Burkholderia* (Okoh et al. 2001), and *Paraburkholderia* (Dias et al. 2019; Lee and Jeon 2018), but these organisms often do not have the enzymes to run the complete metabolic pathways for the degradation of all hydrocarbons present in diesel fuel, especially the aromatic constituents. For example, in a study of rhizoremediation of diesel-contaminated soils, a scarcity of ring-hydroxylating and ring-cleavage dioxygenases among *Gammaproteobacteria* was reported by Garrido-Sanz et al. (2019). These researchers also noted that none of the *nahA* genes in the metagenome was affiliated to *Pseudomonas* or even to the *Gammaproteobacteria* class that dominated the PAH-degrading consortium. In contrast, the consortium reported here contains the CDSs required for the complete degradation of these aromatic components in diesel fuel.

The comparison made between the MAGs assembled from the metagenome data of the enrichment culture and those obtained from studies of similar sites (Eze et al. 2020) revealed the relatively higher abundance, in the consortium, of genes involved in hydrocarbon degradation. These include the *adhP* and *yiaY* genes encoding alcohol dehydrogenases (Drewke and Ciriacy 1988; Glasner et al. 1995; Williamson and Paquin 1987), and the *cpnA* and *chnB* involved in the degradation of cycloalkanes (Iwaki et al. 2002; Sheng et al. 2001). These genes are also involved in the degradation of other organic contaminants such as haloalkanes (Belkin 1992; Yokota et al. 1986). This difference in potential degradative capacity between the MAGs from the two studies can be explained by the taxonomic differences between the MAGs obtained in both cases. In the study of a crude oil bore hole (Eze et al. 2020), majority of the reconstructed MAGs were affiliated to *Gammaproteobacteria*. In contrast, *Alphaproteobacteria*, especially *Acidocella* was the dominant genus in both the enrichment culture and the MAGs from the enrichment culture.

The potential of the enrichment culture to degrade recalcitrant hydrocarbons was also revealed by the presence of genes encoding enzymes involved in degradation of recalcitrant organic compounds. For example, one of the three MAGs from the enrichment culture contained genes encoding 2-halobenzoate 1,2-dioxygenase (*cbdA*), an enzyme that activates the oxidation of 2-chlorobenzoate to catechol. In contrast, none of the 36 MAGs from the previous study contains this gene. Since the enrichment culture is composed of predominantly *Acidocella* strains, the abundance of genes that putatively encode for the degradation of cycloalkanes and aromatic hydrocarbons in the MAGs classified as *Acidocella* is an indication for the potential of the consortium for petroleum hydrocarbon biodegradation.

## Conclusions

The degradation of petroleum hydrocarbons requires several microorganisms with both the ability to withstand toxicity and to harbour the required metabolic pathways. Therefore, a foremost step in establishing a successful microbially-mediated bioremediation approach is the selective cultivation of a suitable microbial consortium with the required degradative capability for the target contaminants. Through successive enrichment using soil samples taken from an historical oil-contaminated site in Germany, we successfully generated a bacterial consortium capable of degrading diesel fuel. We further reconstructed a total of 18 genomes from both the original soil sample and the isolated consortium. The analysis of both the metagenome of the consortium and the reconstructed metagenome-assembled genomes shows that the most abundant bacterial genus in the consortium, *Acidocella*, possess many of the coding DNA sequences required for the degradation of diesel fuel aromatic hydrocarbons, which are often the most toxic components. This can explain why this genus proliferated in all the enrichment cultures. Therefore, this study revealed that the microbial consortium isolated in this study or its dominant genus, *Acidocella*, could potentially serve as an effective inoculum for biotechnological applications in the reclamation of soils contaminated with diesel fuel.

## Author Contributions

Conceptualization and design: MOE, GCH, SCG and RD. Planning and implementation: MOE and RD. Experiments and bioinformatics analyses: MOE. Writing – original draft: MOE. Writing – review and editing: GCH, SCG and RD. Supervision: GCH, SCG and RD. All authors interpreted the results, and agreed to the final version of the manuscript.

## Acknowledgments

The authors would like to thank Macquarie University and the Commonwealth Government of Australia for supporting this research project by providing M.O.E. with an international Research Training Program (iRTP) scholarship, and the German Academic Exchange Service (DAAD) for providing M.O.E. with a DAAD scholarship (Allocation Numbers: 2017561 and 91731339, respectively). This publication was supported financially by the Open Access Publication Fund of the University of Göttingen. The funders had no role in study design, data collection, and interpretation, or the decision to submit the work for publication. We also thank Dr. Anja Poehlein and Melanie Heinemann for assistance during the sequencing.

## Data Availability

Raw sequencing data has been deposited in the sequence read archive of the National Center for Biotechnology Information under BioProject number **PRJNA612814**.

