## Supplementary Materials for "Diversity and metagenome analysis of a hydrocarbon-degrading bacterial consortium from asphalt lakes located in Wietze, Germany"

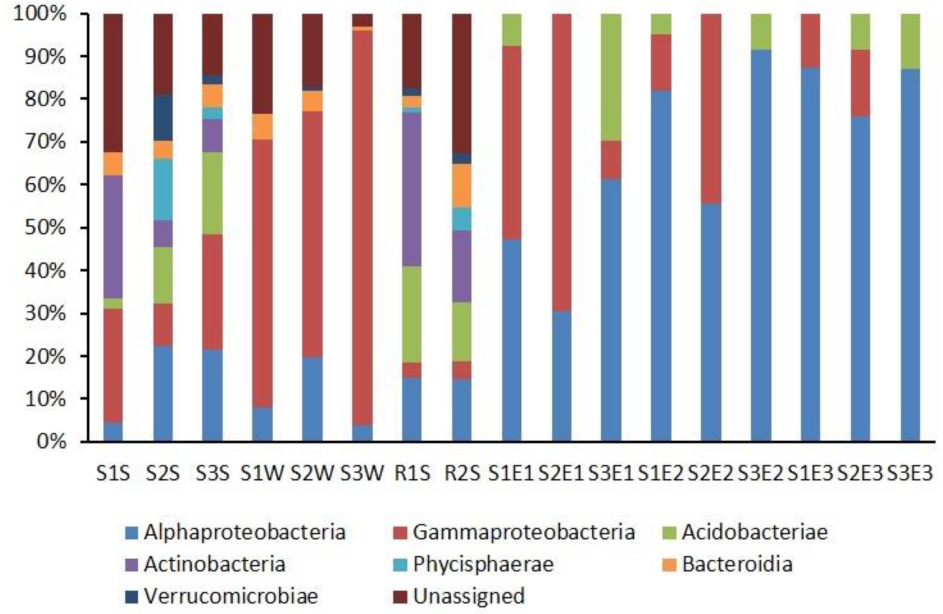


**Supplementary Figure S1.** Bacterial taxonomic distribution of all samples based on 16S rRNA gene amplicon data.

**Supplementary Table S2.** Richness, diversity and evenness obtained from the 16S rRNA sequencing of sampling sites and enrichment cultures

| Sample ID | Name | Richness | Chao1 | Chao1 (%) | Shannon Diversity | Pielou's Evenness |
|---|---|---|---|---|---|---|
| X20200114.BEWE.a.2136_S261 | S1S | 381.99 | 475.51 | 80.33 | 4.140482857 | 0.696418531 |
| X20200114.BEWE.a.2137_S262 | S2S | 958.43 | 1166.75 | 82.15 | 5.927652696 | 0.863422675 |
| X20200114.BEWE.a.2138_S263 | S3S | 885.54 | 1076.78 | 82.24 | 5.625668073 | 0.828986773 |
| X20200114.BEWE.a.2139_S264 | S1W | 731.44 | 971.95 | 75.25 | 4.491982504 | 0.68111784 |
| X20200114.BEWE.a.2140_S265 | S2W | 666.94 | 848.21 | 78.63 | 3.840835511 | 0.590652415 |
| X20200114.BEWE.a.2141_S266 | S3W | 389 | 581.5 | 66.9 | 2.409478261 | 0.40403223 |
| X20200114.BEWE.a.2142_S267 | R1S | 629.32 | 806.44 | 78.04 | 4.89315041 | 0.759258936 |
| X20200114.BEWE.a.2143_S268 | R2S | 1168.08 | 1274.35 | 91.66 | 6.322569919 | 0.89515298 |
| X20200114.BEWE.a.2144_S269 | S1E1 | 62.34 | 67.76 | 92 | 2.684138231 | 0.649503002 |
| X20200114.BEWE.a.2145_S270 | S2E1 | 44.91 | 50.35 | 89.2 | 2.228003222 | 0.585598434 |
| X20200114.BEWE.a.2146_S271 | S3E1 | 38.41 | 40.95 | 93.8 | 1.755745383 | 0.481247923 |
| X20200114.BEWE.a.2147_S272 | S1E2 | 45.09 | 56.52 | 79.78 | 1.896011697 | 0.497815886 |
| X20200114.BEWE.a.2148_S273 | S2E2 | 31.88 | 35.16 | 90.68 | 1.921605494 | 0.555059859 |
| X20200114.BEWE.a.2149_S274 | S3E2 | 14.05 | 14.66 | 95.82 | 0.829529079 | 0.313903749 |
| X20200114.BEWE.a.2150_S275 | S1E3 | 23.26 | 30.09 | 77.3 | 1.761694985 | 0.55984851 |
| X20200114.BEWE.a.2151_S276 | S2E3 | 27.44 | 40.95 | 67 | 1.257543211 | 0.379692792 |
| X20200114.BEWE.a.2152_S277 | S3E3 | 15.28 | 19.03 | 80.3 | 0.962736927 | 0.353097786 |

**Supplementary Table S2**. Metagenome-assembled genomes (MAGs) from both the soil and the enrichment metagenomes.

| User_genome | Source metagenome | classification | closest_placement_taxonomy | closest_placement_ani | aa_percent |
|---|---|---|---|---|---|
| BEWE_m_45_metabat2.1 | Soil sample | f__Acidobacteriaceae;g__Terracidiphilus;s__ | s__Terracidiphilus sp002314435 | 80.06 | 73.89 |
| BEWE_m_45_metabat2.11 | Soil sample | f__Mycobacteriaceae;g__Williamsia_A;s__ | s__Williamsia_A herbipolensis | 80.27 | 89.46 |
| BEWE_m_45_metabat2.15 | Soil sample | f__Acetobacteraceae;g__Acidocella;s__ | N/A | N/A | 74.21 |
| BEWE_m_45_metabat2.18 | Soil sample | f__Steroidobacteraceae;g__;s__ | N/A | N/A | 71.59 |
| BEWE_m_45_metabat2.19 | Soil sample | f__Koribacteraceae;g__Koribacter;s__ | s__Koribacter sp003151155 | 89.00 | 70.85 |
| BEWE_m_45_metabat2.21 | Soil sample | f__Rhodanobacteraceae;g__Rudaea;s__ | s__Rudaea cellulosilytica | 78.03 | 68.95 |
| BEWE_m_45_metabat2.22 | Soil sample | f__UBA5335;g__UBA5335;s__ | s__UBA5335 sp002862435 | 94.10 | 61.21 |
| BEWE_m_45_metabat2.25 | Soil sample | f__UBA5335;g__;s__ | N/A | N/A | 94.40 |
| BEWE_m_45_metabat2.26 | Soil sample | f__Nevskiaceae;g__Solimonas;s__ | N/A | N/A | 65.24 |
| BEWE_m_45_metabat2.27 | Soil sample | f__Burkholderiaceae;g__BOG-994;s__ | N/A | N/A | 86.96 |
| BEWE_m_45_metabat2.28 | Soil sample | f__UBA4822;g__UBA4822;s__ | N/A | N/A | 73.71 |
| BEWE_m_45_metabat2.33 | Soil sample | f__Actinomycetaceae;g__Pauljensenia;s__ | N/A | N/A | 81.75 |
| BEWE_m_45_metabat2.35 | Soil sample | f__UBA5335;g__;s__ | N/A | N/A | 87.62 |
| BEWE_m_45_metabat2.41 | Soil sample | f__Acetobacteraceae;g__;s__ | N/A | N/A | 66.59 |
| BEWE_m_45_metabat2.6 | Soil sample | f__Burkholderiaceae;g__Caballeronia;s__ | N/A | N/A | 93.91 |
| BEWE_m_46_metabat2.2 | Enrichment culture | f__Acetobacteraceae;g__Acidocella;s__ | s__Acidocella aminolytica | 82.29 | 82.02 |
| BEWE_m_46_metabat2.4 | Enrichment culture | f__Acidobacteriaceae;g__Acidobacterium;s__ | s__Acidobacterium capsulatum | 85.13 | 97.02 |
| BEWE_m_46_metabat2.5 | Enrichment culture | f__Acetobacteraceae;g__Acidocella;s__ | N/A | N/A | 96.83 |

**Supplementary Table S3**. Quality check for the MAGs

| Bin.Id | Marker.lineage | X..genomes | X..markers | X..marker.sets | X0 | X1 | X2 | X3 | X4 | X5. | Completeness | Contamination | Strain. Hetero-geneity |
|---|---|---|---|---|---|---|---|---|---|---|---|---|---|
| BEWE_m_45_metabat2.1 | k__Bacteria (UID3187) | 2258 | 187 | 116 | 54 | 125 | 8 | 0 | 0 | 0 | 82.75 | 6.47 | 12.5 |
| BEWE_m_45_metabat2.11 | o__Actinomycetales (UID1814) | 148 | 572 | 276 | 25 | 547 | 0 | 0 | 0 | 0 | 97.53 | 0 | 0 |
| BEWE_m_45_metabat2.15 | o__Rhodospirillales (UID3754) | 63 | 336 | 201 | 48 | 284 | 4 | 0 | 0 | 0 | 93.75 | 1.49 | 25 |
| BEWE_m_45_metabat2.18 | c__Gammaproteobacteria (UID4202) | 67 | 481 | 276 | 110 | 329 | 41 | 0 | 1 | 0 | 77.26 | 8.06 | 0 |
| BEWE_m_45_metabat2.19 | k__Bacteria (UID3187) | 2258 | 188 | 117 | 55 | 132 | 1 | 0 | 0 | 0 | 87.39 | 0.85 | 0 |
| BEWE_m_45_metabat2.21 | f__Xanthomonadaceae (UID4214) | 55 | 659 | 290 | 103 | 536 | 20 | 0 | 0 | 0 | 91.72 | 3.88 | 30 |
| BEWE_m_45_metabat2.22 | c__Gammaproteobacteria (UID4201) | 1164 | 275 | 174 | 93 | 179 | 3 | 0 | 0 | 0 | 64.27 | 1.72 | 66.67 |
| BEWE_m_45_metabat2.25 | c__Gammaproteobacteria (UID4267) | 119 | 544 | 284 | 28 | 513 | 3 | 0 | 0 | 0 | 94.55 | 0.82 | 0 |
| BEWE_m_45_metabat2.26 | k__Bacteria (UID203) | 5449 | 104 | 58 | 48 | 52 | 4 | 0 | 0 | 0 | 66.07 | 6.03 | 50 |
| BEWE_m_45_metabat2.27 | c__Betaproteobacteria (UID3888) | 323 | 387 | 234 | 41 | 340 | 5 | 1 | 0 | 0 | 89.01 | 0.88 | 25 |
| BEWE_m_45_metabat2.28 | k__Bacteria (UID203) | 5449 | 102 | 56 | 13 | 89 | 0 | 0 | 0 | 0 | 80.36 | 0 | 0 |
| BEWE_m_45_metabat2.33 | f__Actinomycetaceae (UID1531) | 42 | 420 | 211 | 58 | 358 | 2 | 2 | 0 | 0 | 87.1 | 2.37 | 25 |
| BEWE_m_45_metabat2.35 | c__Gammaproteobacteria (UID4267) | 119 | 544 | 284 | 44 | 493 | 7 | 0 | 0 | 0 | 90.77 | 1.88 | 14.29 |
| BEWE_m_45_metabat2.41 | o__Rhodospirillales (UID3754) | 63 | 336 | 201 | 114 | 219 | 3 | 0 | 0 | 0 | 65.49 | 0.76 | 66.67 |
| BEWE_m_45_metabat2.6 | g__Burkholderia (UID4006) | 64 | 769 | 248 | 51 | 692 | 26 | 0 | 0 | 0 | 94.03 | 1.51 | 38.46 |
| BEWE_m_46_metabat2.2 | o__Rhodospirillales (UID3754) | 63 | 336 | 201 | 80 | 256 | 0 | 0 | 0 | 0 | 72.39 | 0 | 0 |
| BEWE_m_46_metabat2.4 | k__Bacteria (UID3187) | 2258 | 188 | 117 | 1 | 186 | 1 | 0 | 0 | 0 | 99.79 | 0.85 | 0 |
| BEWE_m_46_metabat2.5 | o__Rhodospirillales (UID3754) | 63 | 336 | 201 | 0 | 335 | 1 | 0 | 0 | 0 | 100 | 0.08 | 0 |
